# The structural landscape of the immunoglobulin fold by large-scale *de novo* design

**DOI:** 10.1101/2023.10.03.560637

**Authors:** Jorge Roel-Touris, Lourdes Carcelén, Enrique Marcos

## Abstract

*De novo* designing immunoglobulin-like frameworks that allow for functional loop diversification shows great potential for crafting antibody-like scaffolds with fully customizable structures and functions. In this work, we combined *de novo* parametric design with deep-learning methods for protein structure prediction and design to explore the structural landscape of 7-stranded immunoglobulin domains. After screening folding of nearly 4 million designs, we have assembled a structurally diverse library of ∼50,000 immunoglobulin domains with high-confidence AlphaFold2 predictions and structures diverging from naturally occurring ones. The designed dataset enabled us to identify structural requirements for the correct folding of immunoglobulin domains, shed light on β-sheet-β-sheet rotational preferences and how these are linked to functional properties. Our approach eliminates the need for preset loop conformations and opens the route to large-scale *de novo* design of immunoglobulin-like frameworks.

## INTRODUCTION

Immunoglobulin-like (Ig) domains are the structural frameworks within the variable region of antibodies, providing anchorage for hypervariable loops engaged in molecular recognition. Despite the widespread use of monoclonal antibodies as protein therapeutics, imaging agents or affinity reagents for biological research, there has been considerable interest in developing antibody-like scaffolds^1–4^ with improved biophysical properties and more programmable structures. *De novo* designing protein frameworks amenable for functional loop diversification, and without relying on naturally existing proteins, holds promise for crafting antibody-like scaffolds with more tunable structures (and functions), alongside the excellent biophysical properties associated with *de novo* designed proteins^5,6^.

Ig domains in nature are β-sandwiches containing between 7 and 9 β-strands arranged in two antiparallel β-sheets packing face-to-face^7,8^. Ideally, their overall geometry can be described by six geometrical transformations describing the relative orientation of two flat β-sheets: three rotations (one twist and two tilts) and three translations along three orthogonal axes passing through the center of the β-sandwich. One notable structural feature of the Ig fold is the arrangement of β-sheets by means of β-strand connecting loops, as they play a crucial role in determining the overall shape of the domain. In previous studies^9,10^, we found that the conformation of β-arch loops^11^, which are crossover connections between β-strands from opposing β-sheets, is key in the formation of β-sandwiches and establishment of the relative orientation of both β-sheets. Specifically, certain combinations of β-arch loop conformations, when compatible with β-strand length and sidechain orientations, strongly support the formation of the central cross-β motif within the core of Ig domains^10^. Based on these principles, we succeeded to *de novo* design 7-stranded Ig domains by a fragment-based computational approach using specific combinations of β-arch conformations^10^. However, the diversity of the designs was heavily constrained by the combinations of β-arch loop conformations considered, which exponentially grows with the number loop conformations per loop site. The relative twist rotation between β-sheets is another critical structural feature of Ig domains, as it modulates the spatial arrangement of the top and bottom faces of the β-sandwich structure, which constitute hotspots for functional binding loops, as seen in complementarity-determining regions (CDRs) of antibodies. Understanding these structural features of the Ig fold is essential for designing Ig-like proteins with tailored functionalities.

Towards fully controllable *de novo* design of Ig domains, we have developed a parametric approach to systematically explore the structural landscape of Ig domains through diversity in their β-strand connections, thus eliminating the need for preset loop conformations. By combining physics-based protein design with deep learning (DL) methods for structure prediction and design^12^, we generated an extensive library of novel Ig domains with highly confident, accurate and convergent AlphaFold2^13^ (AF2) predictions. Our *de novo* immunoglobulins belong to the same fold as naturally occurring Ig domains found in antibodies or nanobodies, while exploring a more extensive structural space. We delved into the diversity of β-arch conformations in our designs, uncovering loop configurations controlling the Ig twist rotation, and hence the spatial organization of potential functionalization sites. Furthermore, we investigated the capabilities of different DL-based methods for sequence design and protein backbone generation for rescuing and building *de novo* Ig scaffolds. In summary, our study offers a comprehensive exploration of the structural landscape of the Ig fold in the realm of *de novo* protein design.

## RESULTS

### Parametric *de novo* design of immunoglobulin domains from ideal β-sheets

We set out to *de novo* design 7-stranded Ig domains through systematic exploration of the β-sandwich conformational space by a parametric approach (Figure 1). We precomputed a library of ideal β-sheets, formed by 3 or 4 antiparallel β-strands of varying residue length (6-8 residues per β-strand) and having optimal backbone hydrogen bond pairing without register shift (Figure S1). Pairs of 3- and 4-stranded β-sheets were then combined through four geometrical transformations describing the overall geometry of the immunoglobulin β-sandwich: the twist rotation around an axis connecting the center of the two β-sheets, the translation along the same axis and translations along the two other orthogonal axes. We finely sampled twist rotations and translations between -60º and +60º and from 10 to 12 Å, respectively. The generated backbones were then closed according to the Ig topology (Figure 1A) by fragment-based design of β-hairpin (connecting two paired β-strands) and β-arch (connecting two unpaired β-strands and crossing over the β-sandwich) loops; resulting in fully connected Ig backbones. Fundamental rules have been described for designing β-hairpin^14^ and β-arch^9^ loops based on the orientation of their two neighboring sidechains (Figure 1B (2), inset). We designed canonical β-hairpins with 2 and 5 residues for L and R chiralities, respectively^15^. For β-arches, instead, we explored loop lengths between 3 and 6 residues, for each of the four possible sidechain directionality patterns, without any restriction in their conformation. We first bridged β-strands E5 and E6 (loop L5 in Figure 1A) and then connected β-strands E2 to E3 and E4 to E5 simultaneously (loops L2 and L4 in Figure 1A, respectively) to ensure compatibility – L2 and L4 are adjacent in the Ig structure. For all closed backbones, we designed 5 amino acid sequences with Rosetta^16^ sidechain packing calculations, and probed the capability of the designed sequences to recapitulate their structures by AF2 protein structure prediction (without multiple sequence alignment or template information) (Figure 1A).

**Figure 1.**
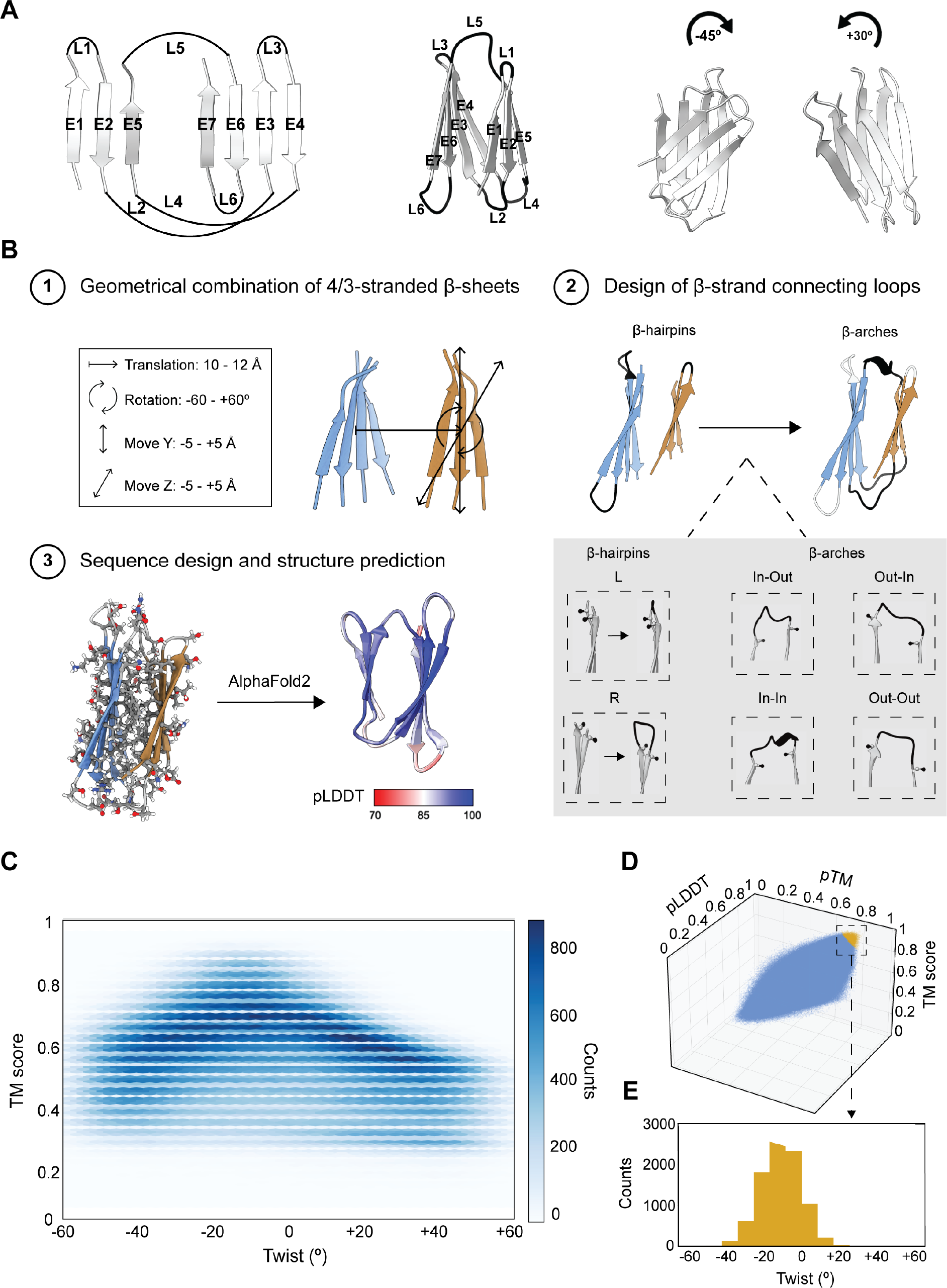
Design and characterization of immunoglobulin-like domains. **A**, The immunoglobulin structure is composed by 2 β-sheets packing face-to-face. The 7 β-strands (E1-7) are connected through 3 β-hairpins (L1, L3 and L6) and 3 β-arches (L2, L4 and L5). Clockwise relative twists of the 3-stranded β-sheet are considered negative rotations, while anticlockwise twists are considered positive rotations. **B**, Workflow for the *de novo* design of Ig domains from ideal β-sheets: 1) Pairs of 3- and 4-stranded β-sheets are combined through a series of geometrical transformations. 2) The design of canonical β-hairpins is followed by the design of structurally unconstrained β-arches (L5 followed by L2 and L4 simultaneously). Bottom inset shows the different chiralities considered for both types of ββ loops. 3) For each closed scaffold, 5 different sequences were designed and subsequently validated by AlphaFold2 protein structure predictions. **C**, *In silico* validation of the nearly 2.5 million *de novo* designed immunoglobulins as a function of the average TM score of the predictions with respect to the β-sheet-β-sheet twist. Darker areas represent more populated regions. **D**, 3D representation of the immunoglobulin designs as a function their averaged pLDDT, pTM and TM scores. Golden points represent highly confident (high pLDDT and pTM scores) and accurate (high TM scores) predictions. **E**, Twist rotation distribution for 8,942 high-quality designs.

Following this approach, we designed ∼2.5 million *de novo* Ig scaffolds, ensuring that all different combinations of variables (geometrical parameters, β-sheet length combinations and loop lengths) were uniformly sampled (Figure S2). After protein structure prediction, we examined the distributions of the sampled twist rotations as a function of their similarity to the AF2 predicted models. While the whole range of rotations is sampled, more density is found around the negative twist (Figure 1C, the darker the denser). For the *in silico* validations with AF2, we classify a design as high-quality if (1) average RMSD ≤ 1.25 Å (across the top-3 AF2 models), (2) standard deviation of RMSD ≤ 0.25 Å (considering all 5 AF2 predictions) and (3) average composite score^17^ (TM-score^18^ * pTM * pLDDT) ≥ 0.5 (across the top-3 AF2 models). By applying this selection criteria, we ensure that the selected Igs closely recapitulate the structure of the design, while having highly confident and convergent AF2 predictions. Convergence across predictions has been related to success for *de novo* designed proteins^19^. From the pool of designs, we have identified close to 9,000 Ig domains with excellent predictions, whose composite scores range from 0.5 to 0.74 (Figure 1D, yellow dots); thus representing very high confident yet accurate predictions. In terms of twist rotations, the distribution of these designs is also skewed to negative values with a median of -11.3º (Figure 1E), sampling the whole range of β-sheet-β-sheet translations (median of 10.9 Å) (Figure S3). In fact, out of the 8,942 scaffolds, only 1,259 (14%) have positive rotations with a median of +4.0º and a maximum rotation of +28.7º.

### Improving *de novo* immunoglobulins with deep learning

We reasoned that the cause for medium-to-low quality structure prediction of many designs could arise from (1) faulty amino acid sequences installed in good backbones, or (2) deficient β-strand-β-strand connecting loops. We assessed whether two advanced DL-based design methods could address these two problems and hence rescue the designs: fixed backbone sequence design with ProteinMPNN^20^, and joint construction of sequence and structure of ββ loops by inpainting as implemented in RFDesign^21^. For sequence-redesign with ProteinMPNN, we selected 2,891 designs with high confidence and accurate predictions (composite score ≥ 0.5) but low structural convergence across the five AF2 models (Figure 2A); suggesting that the designed backbone is reachable but not strongly encoded by the sequence. For inpainting, we selected 67,302 designs with moderate-to-severe structural deviations in the prediction of the loops connecting β-strands, regardless of their confidence metrics or structural convergence (Figure 2B). For these designs, those loops showing such predicted deviations were entirely rebuilt by inpainting (from 1 to 6 depending on the design) using RFDesign.

**Figure 2.**
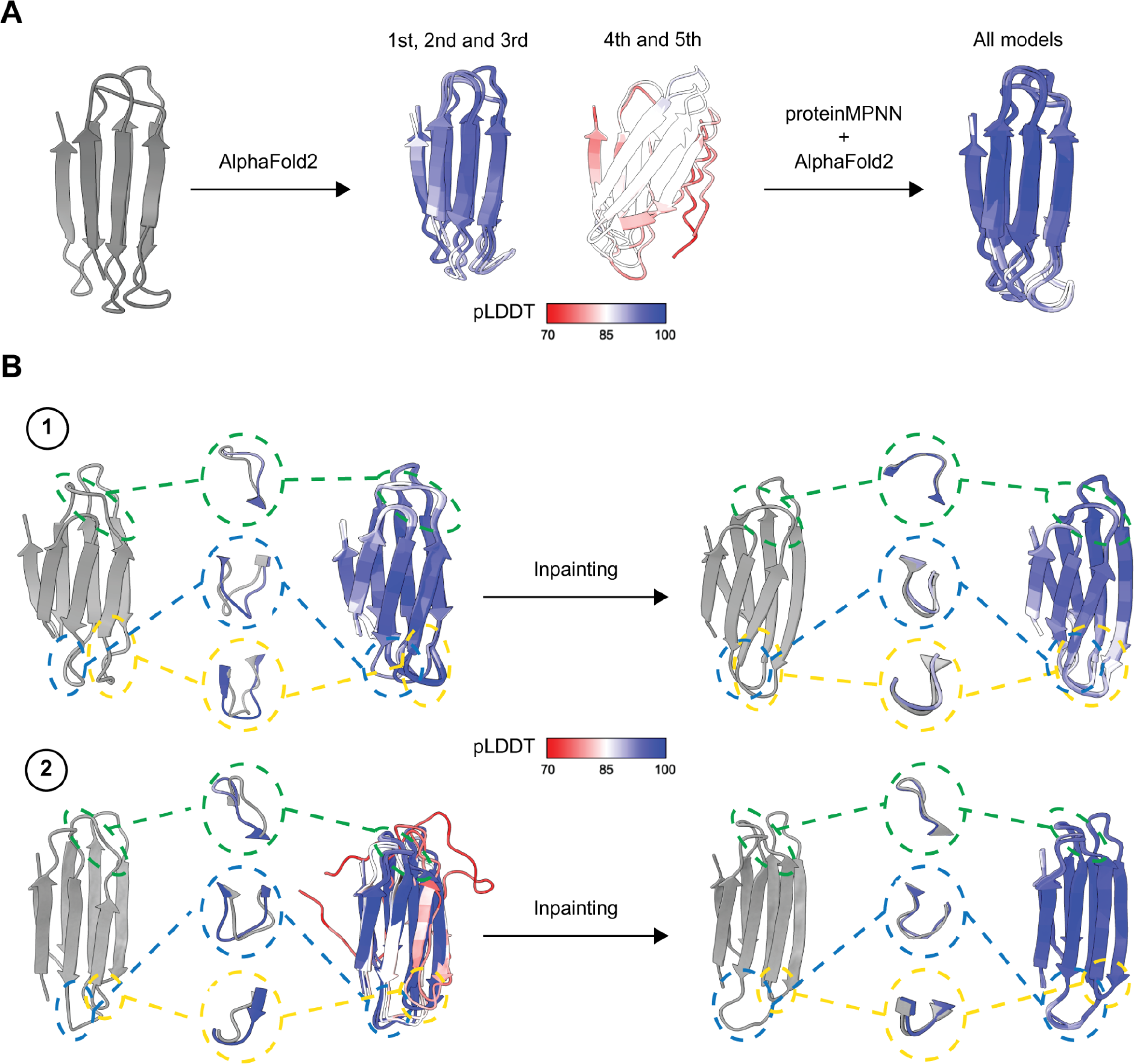
Rescue of inaccurate immunoglobulin scaffolds by deep learning. **A**, Scaffolds with accurate yet non-convergent protein structure predictions were selected for sequence redesign with ProteinMPNN and subsequent validations by AF2. **B**, The two different scenarios considered for structure redesign. 1) Scaffolds with convergent or 2) non-convergent protein structure predictions with severe inaccuracies on the loops were redesigned by inpainting with RFDesign. The resulting models were further validated by AF2 predictions.

The selected scaffolds for sequence-redesign span twist rotations between -53.5º and +46.2º and follow a distribution centered at -21º (Figure 3A). For each of them, we generated 50 sequences with ProteinMPNN (and energy minimized with FastRelax^16^) and predicted their structure with AF2. We found that for 2,492 unique scaffolds (and sequences) the non-convergence issue was resolved, yet having high confidence prediction metrics (composite score up to 0.77). This pool of successful redesigns represents 86% of the total selected designs. After recalculating their β-sheet-β-sheet twist rotations, we found that 75% (1,863) were within the same rotational range (Figure 3B), while 629 (25%) significantly changed rotations (Figure 3C) (as compared to their original designs) to accommodate the newly designed sequence. Similarly, the selected scaffolds for loop structural repair span rotations between -60º and +60º and are centered around 0º (Figure 3D). For each scaffold, we used inpainting to rebuild those loops (backbone and sequence) showing structural disagreement between the design and the AF2 predictions, and subsequently predicted the structure of the inpainted designs with AF2. We found that 51% of them (34,335) had composite scores greater than 0.5 (up to 0.77) with averaged RMSDs below 1.25 Å and convergence across the predicted models. As before, while for the vast majority (26,438) the β-sheet-β-sheet twist rotations were within comparable ranges (Figure 3E), we again found shifted rotations for 23% of them (Figure 3F). Both DL design methods showed a high success ratio in rescuing designs and a preference towards less extreme β-sheet-β-sheet twists.

**Figure 3.**
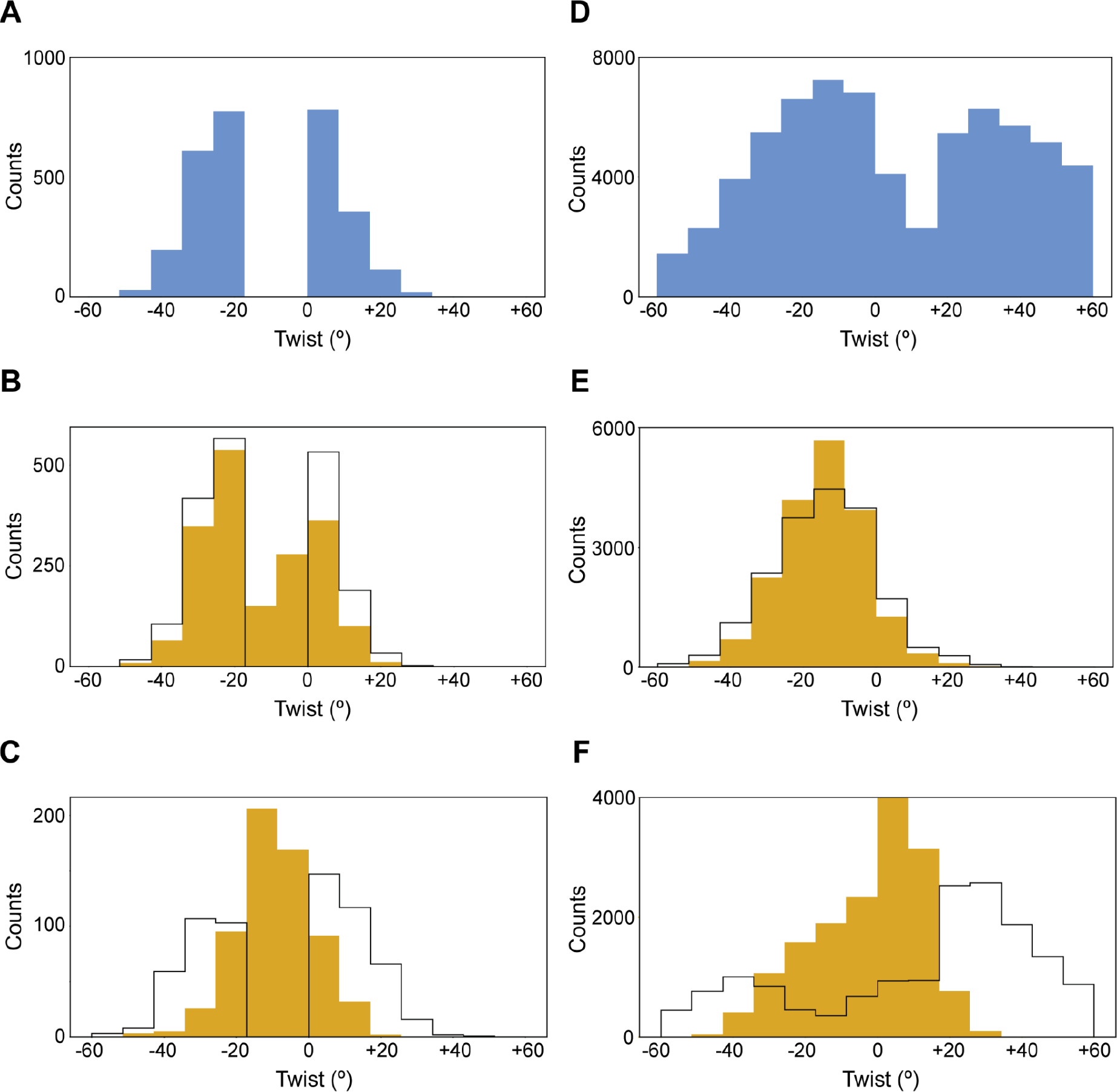
Results for the deep learning-based rescuing protocols. **A**, Selected designs for protein sequence redesign with ProteinMPNN. **B**, Successful sequence-redesigns keeping and changing (**C**) their β-sheet-β-sheet twist with respect to the original designs (**A**). β-sheet-β-sheet twist distributions of original designs (unfilled histograms) and successfully redesigned (yellow histograms). **D**, Selected designs for loop repair with Inpainting. **E**, Successful inpainted designs keeping and changing (**F**) their β-sheet-β-sheet twist with respect to the original designs (**D**). Histograms are displayed as in **B** and **C**. All histograms show the number of counts per bin with respect to the β-sheet-β-sheet relative rotations.

### Comparison of *de novo* immunoglobulins to naturally occurring Fv domains and nanobodies

We pooled all the high-quality designs (as defined above based on AlphaFold2 prediction) coming from the parametric protocol and DL-based rescue, and obtained a total of 45,769 *de novo* Ig domains. This pool of designs spans rotations between -58.5º and +45.7º with a distribution centered around -10º (Figure 4A). We first sought to compare our scaffolds to existing (natural) Ig-like structures. To this end, we compiled and manually curated a dataset of 2,357 non redundant structures including 1,849 antibodies (i.e. Fv, Fab and scFv) and 508 nanobodies from the SAbDab database^22^ (< 70% sequence identity). In order to focus the comparisons on the immunoglobulin domain, we only kept the variable fragments (Fvs) from the Fab regions of all antibody structures. Next, we independently clustered each of the 3 subgroups at a TM-score threshold of 0.8 for the *de novo* immunoglobulins and 0.9 for the Fv domains and nanobodies; resulting in 560, 107 and 91 clusters for designs, Fvs and nanobodies, respectively. For all cluster representatives, we calculated all pairwise maximum TM scores – i.e., without normalization by sequence length – and computed a structural similarity matrix. A clear segregation between our designs and naturally occurring Ig domains can be observed (Figure 4B). In comparison to Fvs and nanobodies, our designs had TM scores ∼0.5 between them, which suggests that all have the same fold while sampling divergent regions of the Ig structural space. For Fvs and nanobodies, we find relatively high TM scores across both groups, underpinning the highly conserved structure of their frameworks responsible for anchoring hypervariable loops. In contrast, the designs sampled a much larger structural diversity, while in the same fold (Figure 4B,C), through our exhaustive exploration of conformational parameters; including β-sandwich geometries, ββ connections and β-sheet sizes.

**Figure 4.**
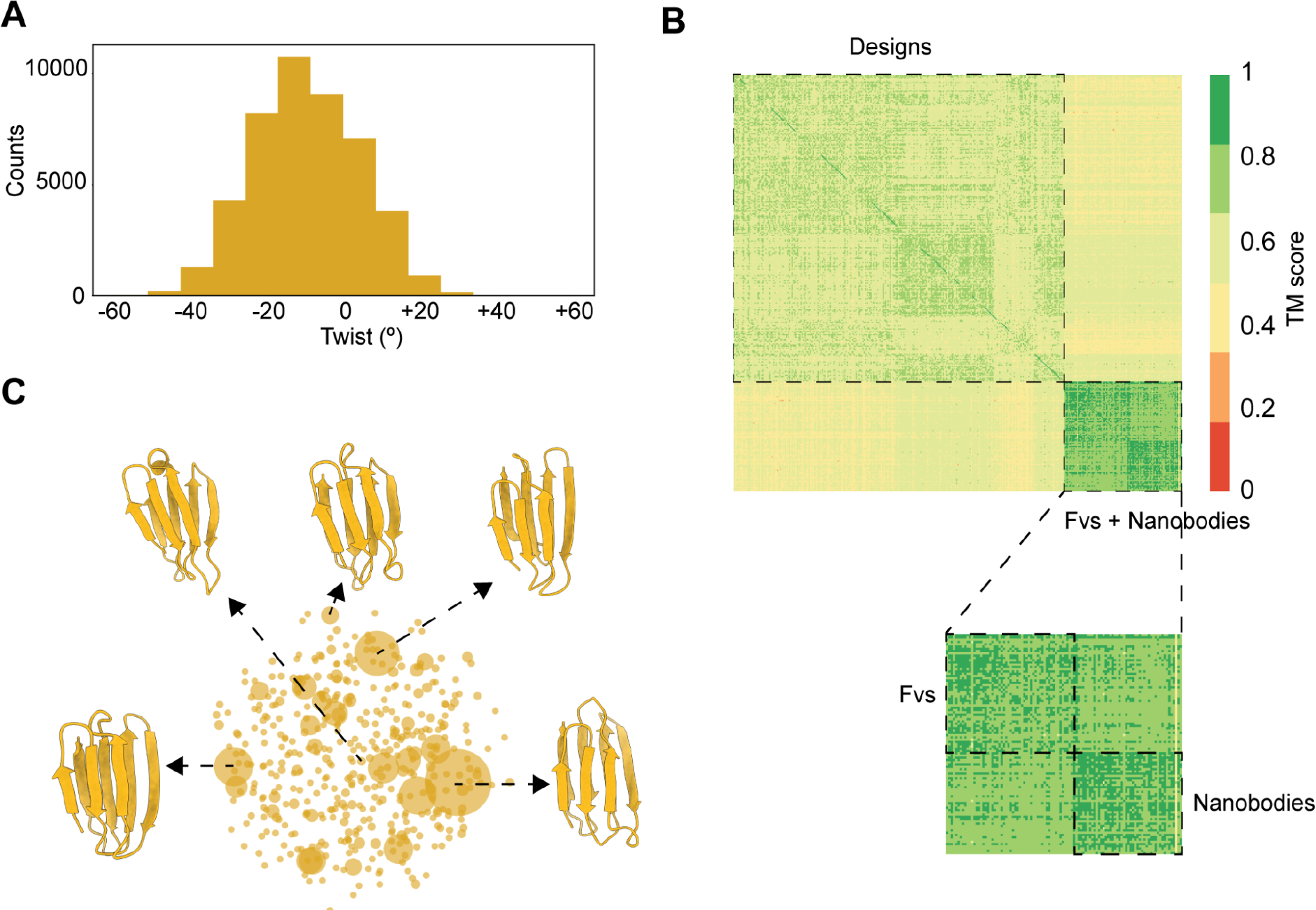
Structural comparison of *de novo* designed Ig scaffolds to natural Ig domains. **A**, Twist rotation distribution of the selected 45,769 high-quality de novo immunoglobulins. **B**, Structural similarity matrix sorted according to the nature of the scaffold: Designs/Fvs/nanobodies. Bottom inset zooms into the Fvs-nanobodies comparison. **C**, Network-like representation of the 560 design clusters originated from **A**. Cartoons show several models from most populated clusters as well as the center of the network.

### Analysis of most frequent β-arches in *de novo* immunoglobulins

We next examined the loop conformations emerging in the pool of high-quality designs. The two face-to-face β-sheets in the immunoglobulin fold are bridged by three β-arch loops, which connect β-strands E2-E3, E4-E5 and E5-E6 (L2, L4 and L5 in Figure 5A, respectively). In contrast to our previous study^10^, here we generated *de novo* Ig domains through diversity in their β-arch loops without constraints in their conformation (or ABEGO type) and up to six amino acids in length. ABEGO backbone torsion bins provide a convenient way to classify the backbone geometry of protein residues based on the Ramachandran plot region of their ϕ/Ψ dihedrals^15^. For each of the high-quality designs, we computed blueprint files and extracted the ABEGO types of the three β-arch loops as defined by DSSP^23^. In the set of nearly 50,000 selected designs, we find a sizable diversity of β-arch conformations with 1,532, 2,132 and 1,371 unique ABEGO types for loops L2, L4 and L5, respectively. This diversity is even larger when considering paired β-arches (16,382, 19,910 and 20,184 for L2:L4, L2:L5 and L4:L5) or the three of them (37,094 for L2:L4:L5).

**Figure 5.**
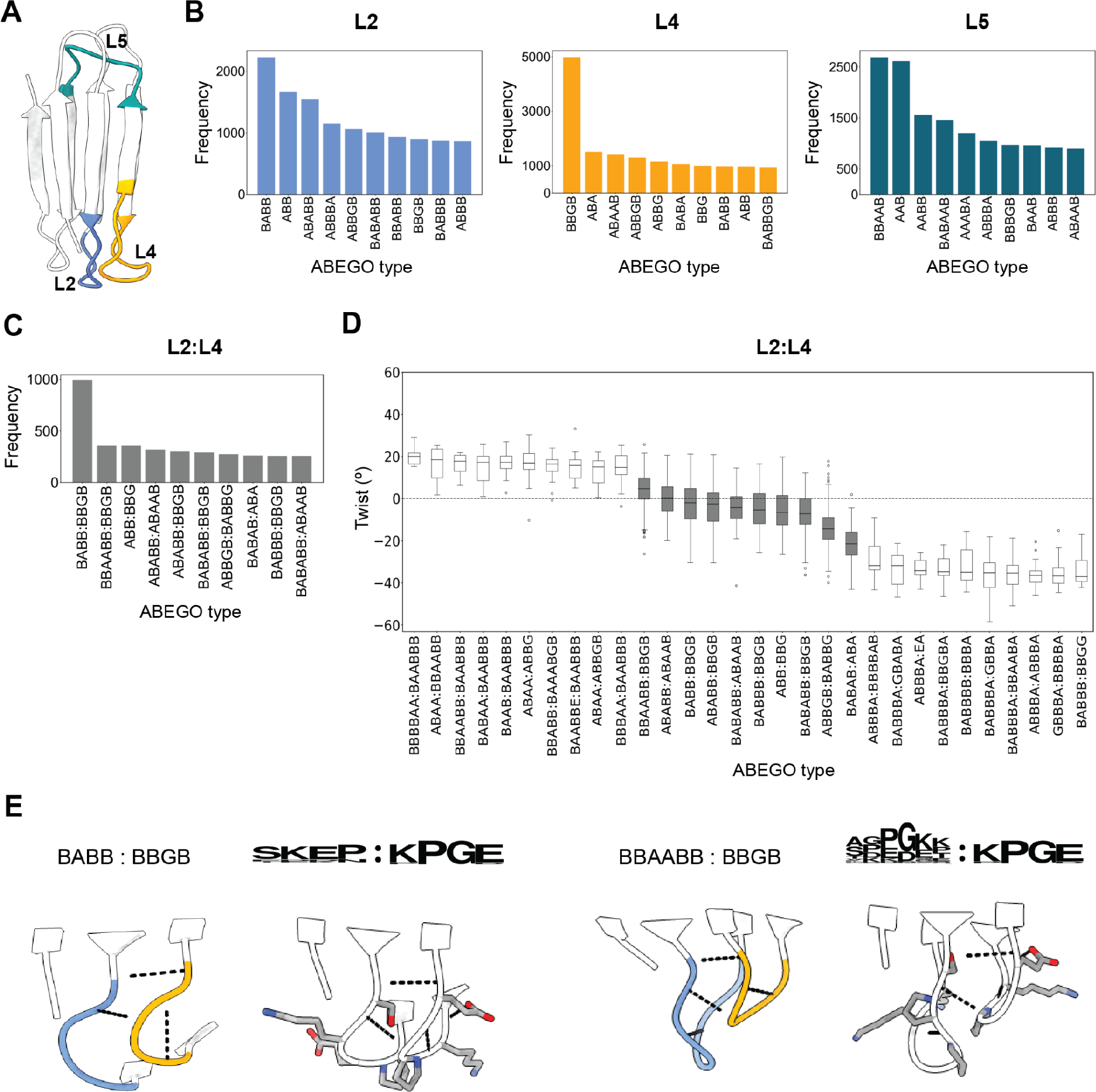
Predominant β-arch configurations found in *de novo* designed immunoglobulins. **A**, The three different β-arches of an immunoglobulin domain. **B**, Most frequently found ABEGO types in our designs for each of the β-arches. **C**, More frequent combinations of ABEGO pairs for the L2:L4 duo. **D**, Distribution of L2:L4 β-arch pairs as a function of the immunoglobulin twist. Gray boxes are the pairs as in **C. E**, Two most frequently found L2:L4 pairs with their sequence logos. Bottom cartoons show interatomic contacts.

However, we observe that certain ABEGOs are more frequently found either in the three β-arches than others (Figure 5B). For example, BBGB is the predominant configuration found in L4, which is also among the most frequently found ABEGO types in L2. Indeed, β-arches L2 and L4 are structurally adjacent in the Ig fold and therefore restrict the configuration of one another. We note this coupling by the increased number of frequently found paired L2:L4 conformations (Figure 5C), which in turn explains the relatively decreased diversity in terms of the number of unique L2:L4 ABEGO pairs as compared to those in L2:L5 and L4:L5. In fact, the predominant L2:L4 paired conformation (BABB:BBGB) is five and two times more frequent than BABB:BBAAB and BBGB:AAB, which are the predominant ABEGOs for L2:L5 and L4:L5, respectively (Figure S4). These observations suggest a stronger cooperation between β-arches L2 and L4, relative to the other two pairs, in stabilizing the structure of the Ig fold.

Besides loop frequencies, we also analyzed whether these non-local ββ connections control the overall geometry of the Ig fold. At a first glance, we noticed that particular loop configurations (alone or in combination) are not shared between positive and negative rotations. For each β-arch, multiple loop configurations were exclusively found in negative twist rotations, while others predominantly favored positive rotations (Figure S5) – less diversity of loops exclusively found in positive rotations was observed. This effect is more evident for pairs of β-arches. While the most frequently found loop:loop conformations shape neutral rotations (between -10º and 10º), other less represented pairs are specific to positive (>10º) or negative twist rotations (<-10º) (Figure 5D and Figure S6). Interestingly, these observations also apply for the L2:L4:L5 trio; where, for example, ABAA:BBGGGBB:AABB and BAAB:BBGB:BAA only shape scaffolds with positive twists of around +15º (Figure S7). Also, the identified β-arches in high-quality Igs sample an immense sequence space. Indeed, we find more than 20,000 unique amino acid sequences for each single loop and more than 30,000 for each loop:loop and loop:loop:loop combination. However, as previously reported, there are certain sequences that preferentially encode for specific loop conformations^9^. As an illustration, for the paired L2:L4 combination, we find 35,885 unique sequences for 16,382 unique ABEGOs. This would roughly indicate that there are, on average, 2 different sequences per loop configuration. However, for the predominant BABB:BBGB configuration, there is one prevalent paired sequence (SKEP:KPGE), which is found in 10% of the cases (Figure 5E). Consistent with our previous observations, we find a higher number of unique sequences in L2:L5 and L4:L5 pairs (41,485 and 44,329), which again suggests a lower cooperativity effect between these β-arch pairs in comparison to that of L2:L4.

### Relative β-sheet-β-sheet rotations control the localization of functionalization spots

The six ββ loops of the Ig fold are spatially clustered into two different groups. Three loops (L1, L3 and L5) on the top side and other three (L2, L4 and L6) on the bottom side (Figure 6A). These two faces represent putative sites where designed ββ loops could collectively build functional paratopes binding targets of interest, hence mimicking the action of paratopes observed in antibodies and nanobodies. We have previously shown that β-hairpin regions in monomeric^10^ and single-chain dimeric^24^ Igs are sweet spots for functional ligand-binding loops, and that their relative orientation could be finely tuned through structural diversification of Ig scaffolds. Here, we analyze the diversity of the ∼50,000 high-quality Ig designs in terms of their loops’ relative orientations in relation to the Ig twist rotation, which highly controls the overall scaffold structure. To this end, we computed the pairwise center-of-mass euclidean distances within each of the two subsets of loops, for each of the selected designs. As a general trend, we observe that a positive twist tends to enlarge the distance between loops on the same side, while negative rotations place them in closer proximity (Figure 6B and Figure S8). The loop-loop distance separation was found to range between 9 Å and 27 Å when twisting from negative to positive angles (Figure 6C); suggesting that negative rotations would enable more interaction (and hence cooperativity) between loops; as often observed in complementarity-determining regions in antibodies. Overall, the immunoglobulin twist controls the global organization of anchoring sites for functional loops, which is key for designing target-tailored scaffolds.

**Figure 6.**
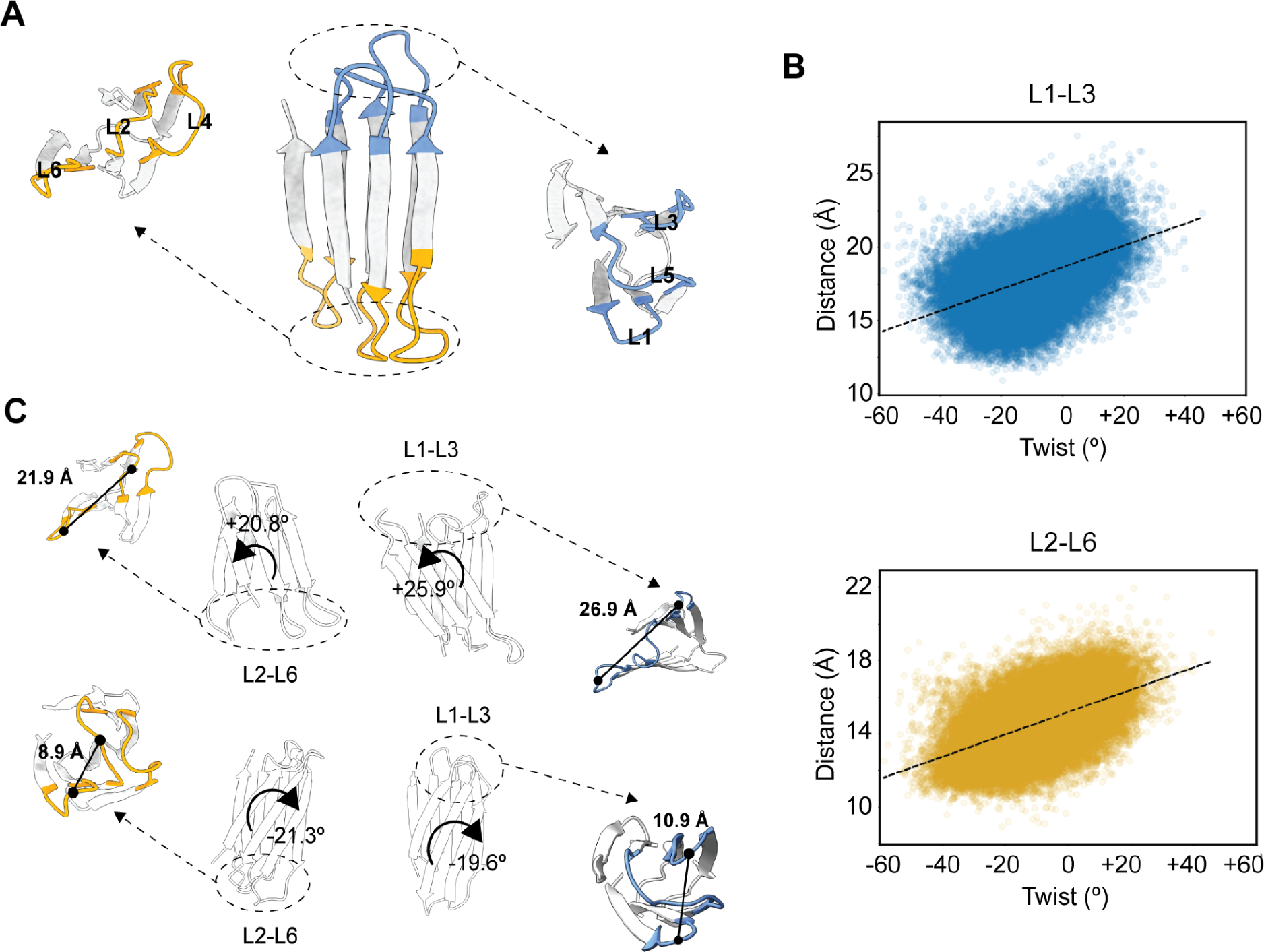
Analysis of pairwise relative distances between putative functionalization spots. **A**, The immunoglobulin fold has 3 functionalization spots on the top face (L1, L3 and L5 in blue) and 3 on the bottom face (L2, L4 and L6 in yellow) **B**, Distribution of distances between the center-of-masses of L1:L3 and L2:L6 (y axis) as a function of the β-sheet-β-sheet twist (x axis). **C**, Examples illustrating how twisting from positive (top) to negative (bottom) rotations impact the relative euclidean distance between functionalization spots.

### Immunoglobulins with antinatural positive twists are harder to design

Out of the 45,769 high-quality immunoglobulins, 47% of the designs are shaped by rather neutral twists ranging from -10º to +10º (21,508), 45% have negative twists from -58.5º to -10.0º (21,508) and the rest (3,567) have twists higher than +10º, representing the latter only the 8% of the total population. We attempted to elucidate the origin of this imbalance by computing several physics-based metrics including the Rosetta (per residue) energy score, buried and accessible surface areas as well as the shape complementary between opposing β-sheets. In general, we notice that immunoglobulins with negative twists tend to be larger (in number of residues) than those with positive ones, which translates into differences in their buried surface areas (BSAs) (Figure 7A) – BSA is closely related with the stability of the native conformation in monomeric formats^25^. Other design scores related to packing efficiency and protein folding stability, such as the Rosetta energy (Figure 7B), also tended to be more favorable for Ig designs with negative twists (Figure S9). These results suggest that non-positive twist Ig scaffolds tend to be better predicted and scored.

**Figure 7.**
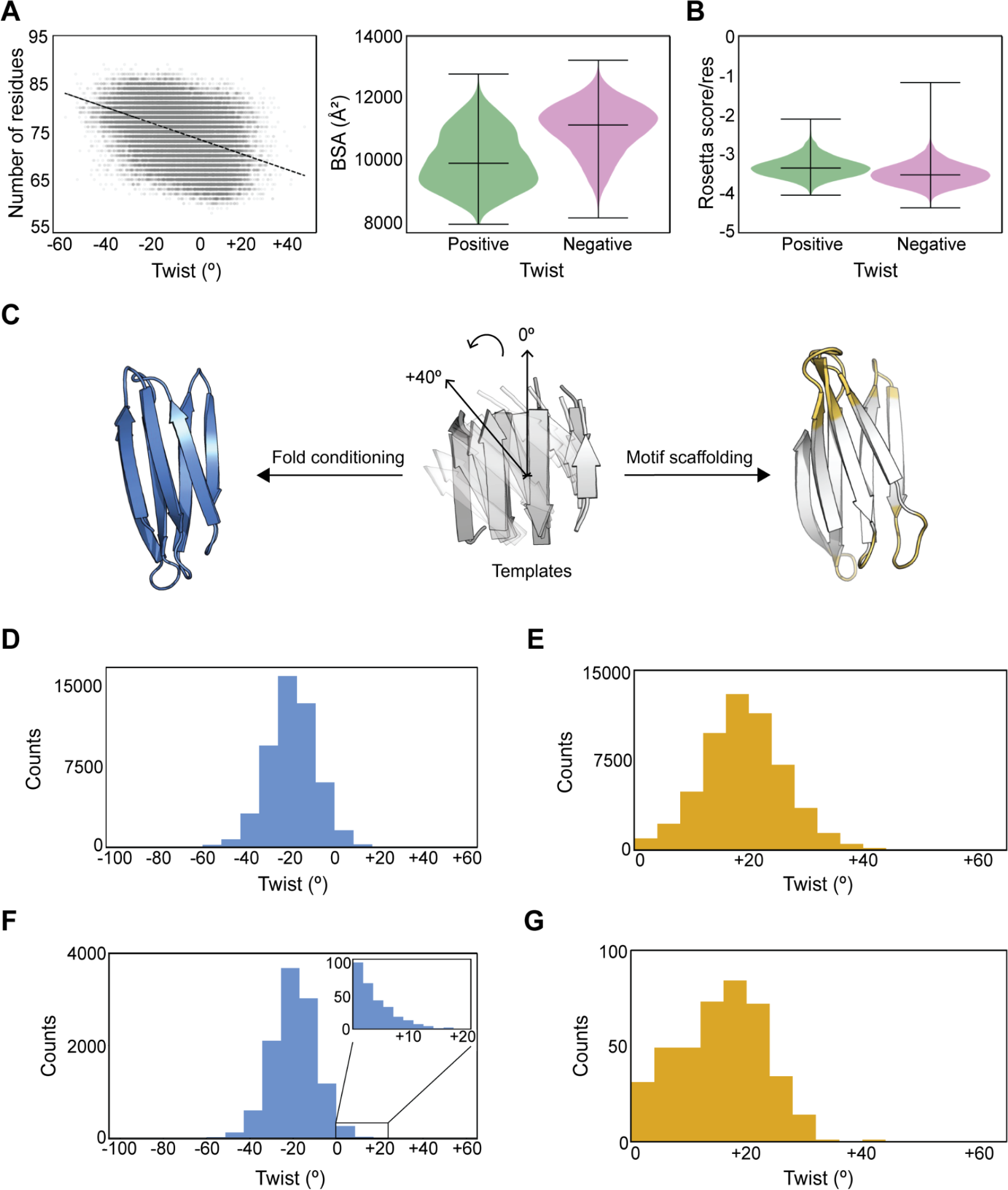
Analysis of immunoglobulin designs with positive twists. **A**, Size (left) and buried surface area (right) distributions of the 45,769 high-quality Ig designs with respect to their β-sheet-β-sheet twist. **B**, Comparison of Rosetta per residue scores (the more negative, the better). **C**, Design of Igs domains from high-quality templates by *fold conditioning* (left) and *motif scaffolding* (right) by RFdiffusion. For the *motif scaffolding* protocol, only the six ββ loops were redesigned for each of the 100 templates, while designs from the *fold conditioning* protocol are newly designed scaffolds. **D**, 50,274 Igs domains generated by the *fold conditioning* protocol with respect to their twist. **E**, 54,897 Igs domains generated by the *motif scaffolding* protocol with respect to their twist. **F**, The 10,690 high-quality scaffolds passing our filtering criteria from the *fold conditioning* protocol. Top-right inset shows the tiny fraction (290 designs) exhibiting positive twists. **G**, The 408 high-quality scaffolds with positive twists passing our filtering criteria from the *motif scaffolding* protocol.

Based on the difficulty of designing Ig domains with positive twists using different protocols, we next set out to design Ig scaffolds with positive twists using RoseTTAFold Diffusion^26^ (RFdiffusion), which has shown outstanding results for *de novo* protein backbone generation and higher performance than other methods in several design tasks. To this end, we designed 200,000 Ig scaffolds by two different protocols included within RFdiffusion: *fold conditioning* and *motif scaffolding* (100,000 designs each). These are two template-based protocols that bias the diffusion model and guide the generation of backbone structures using precomputed structural information. On the one hand, *fold conditioning* extracts secondary structure information and contacts from a given protein structure template to bias the generation of structural diversity around the template fold (Figure 7C, left). On the other hand, *motif scaffolding* allows to build protein segments around a given structural motif (Figure 7C, right). For both methodologies, we used the same template set of 100 high-quality *de novo* 7-stranded immunoglobulin scaffolds with exclusive positive twist rotations ranging from 0º to +40º (Figure 7C, central scheme). While in *fold conditioning* all the structure was built anew, in *motif scaffolding* we rebuilt both β-hairpin and β-arch loop connections using β-sheet residues as the motif. After backbone generation by RFdiffusion, amino acid sequences were designed with ProteinMPNN and validated by protein structure predictions with ESMFold^27^ and AF2 (see Methods for details).

From the pool of 200,000 diffused models, we could unequivocally identify ∼50% of them (50,274 and 54,897 for *fold conditioning* and *motif scaffolding*, respectively. Figure 7D,E) as 7-stranded immunoglobulin domains: 2 well-defined β-sheets (i.e. formed by continuous and regular β-strands) with 3 and 4 antiparallel β-strands, proper strand connectivity and optimal hydrogen backbone pairing, as in our *de novo* designed scaffolds. The remaining designs recapitulate either less regular Igs or misfolded entities (Figure S10). For the *fold conditioning* protocol, 10,980 unique designs showed excellent prediction metrics (as above described), with twists ranging from -81º to +16º, and a distribution centered at -19.4º (Figure 7F). However, only 290 of these high-quality designs exhibit positive twists (2.5%, Figure 7F, inset). For the *motif scaffolding* protocol, instead, a much reduced fraction of the diffused scaffolds had high-quality AF2 predictions (408 unique Igs, which represent less than 1% of the generated scaffolds) (Figure 7G), with twist rotations ranging from +0.02º to +41º and a median of +16.1º. Overall, even with the use of high-quality template data, we observe again an evident bias towards the design of Ig scaffolds with negative twists, as the proportion of generated (and validated) scaffolds with positive twists by RFdiffusion is minor. In line with this observation, designs rescued by ProteinMPNN or inpainting tended to shift positive twist rotations to more neutral ones for a fraction of the designs. Taken together, these results suggest that Ig domains with positive twist rotations are generally more difficult to design and predict.

## DISCUSSION

*De novo* design of all-β proteins lags behind in comparison to all-α or mixed αβ structures^5,6^. Specifically, *de novo* design of β-sandwiches, such as jelly roll and Ig-like structures, has been historically considered of great difficulty due to their tendency to oligomerize through their exposed strand edges and their slow folding kinetics, which can trap folding intermediates. First successful designs of these scaffolds^9,10^ required precise design rules for guiding the (fragment-based) generation of their backbones towards structures that can be strongly encoded by an amino acid sequence (i.e., designable), which in turn allowed to minimize costly assessments of the folding energy landscape^28^. However, the structural diversity of the designs generated by fragment assembly was in part limited by the requirement of enumerating loop conformations favoring folding. Recent methods^29,30^ for *de novo* protein design have been tested on designing some of these all-β folds, including jelly roll and Ig-like, but the experimental results showed that their design still remains challenging. Here, we have developed a parametric approach that eliminates the need of preset design rules to *de novo* design 7-stranded immunoglobulin domains. In combination with DL-based protein design and structure prediction, we have assembled and analyzed about 50,000 highly accurate, confident and diverse 7-stranded Ig-like scaffolds. This approach can be easily adjusted to design β-sandwich domains of varying number of β-strands and with different β-strand connectivity.

Besides structure validation, AF2 predictions enabled the diagnosis of specific design issues. Using AF2 as a diagnostic tool, we demonstrated how DL-based design algorithms can turn low-quality designs into high-quality ones with accurate, confident and convergent AF2 predictions. ProteinMPNN significantly enhanced the structural convergence of AF2 predictions across the five models, whereas inpainting effectively rectified structural deviations in the loops. Both methods proved highly efficient in rescuing designs, with success rates of 86% for ProteinMPNN and 51% for inpainting. Notably, inpainting represents a remarkable advance in addressing the long-standing problem of loop closure in protein design. Yet, Ig domains with positive rotations were found to be the most challenging geometries for these approaches, including the parametric approach. To further address this issue, we explored the application of RFDiffusion to generate positively rotated backbones using a set of high-quality templates as a bias for fold conditioning and motif scaffolding. Although these approaches were effective in producing backbones that can support sequences approved by AF2, they still showed a bias towards neutral or negative twist rotations. In addition to possible limitations of the design methods, the observed rotational preferences may be related to more effective packing arrangements (Figure 4) and conformational loop preferences found in Ig domains with neutral or negative rotations (Figure S11). The difficulty of designing Ig domains with positive twist rotations also agrees with the observation that naturally occurring Ig domains have predominantly negative rotations (median -33º), as recently reported^10^. Interestingly, the design twist distribution is shifted to more neutral values (Figure 4A) (median -10º), which must also contribute to the new structural space explored by the designs.

The DL revolution in protein structure prediction and design minimizes the need for accurate design rules. While DL-based structure prediction allows to accurately and economically assess the folding of designed sequences, DL-based design methods for backbone generation and sequence design offer efficient design solutions with minimal designer input^12^. In light of this paradigm shift, we have shown in this paper that Ig domains can be designed at large scale –i.e. in terms of number and structural diversity of novel Ig domains with high-quality AF2 predictions – by combining parametric and DL methods, without preset loop conformations^10^ favoring the folding of the Ig core structure. We have performed a large-scale AF2 prediction for ∼4 million Ig designs (∼2.5 million from parametric design; ∼145,000 from ProteinMPNN; 310,000 from Inpainting; and 1 million from RFDiffusion). The comprehensive design dataset of high-quality Ig domains assembled here enabled us to identify distinct conformational preferences of β-arch loops and understand the degree of interdependence they exhibit in the Ig fold. Our analyses also uncovered preferential twist rotations of the Ig β-sandwich similar to that found in naturally occurring immunoglobulins^10^. These findings hold significant importance as they influence the overall structure of Ig domains, and hence the spatial arrangement of anchoring sites for functional loops. It will be exciting in future work to investigate the potential of this high-quality scaffold library to anchor flexible, functional loops for protein-binding applications, either through high-throughput screening of loop libraries^31,32^ or computational design of grafted or entirely new loops. Overall, rather than establishing principles in advance for the design process, it is now feasible to derive post-design principles by gathering extensive datasets through DL structure prediction and design. This ultimately opens a different perspective for advancing our understanding on how proteins can be designed from scratch.

## MATERIALS AND METHODS

### *Parametric* design of immunoglobulin domains

The parametric design protocol was implemented using PyRosetta^33^. We built a library of ideal 3-stranded and 4-stranded β-sheets of varying residue length per strand in two different chiralities, as defined by the directionality of the Cβ of the termini residues. For the first chirality, we generated a single poly-valine β-strand of 8 residues long using ϕ/Ψ values of -140º/+130º, which corresponds to the extended region of the Ramachandran plot. Then, we made a copy of this strand, rotated it 180º and performed a lateral translation of 5 Å. This process was repeated until obtaining 3-stranded and 4-stranded antiparallel β-sheets. For the second chirality, instead, we generated a poly-valine β-strand of 9 residues using the same ϕ/Ψ values and trimmed the first valine; effectively forming a β-strand of 8 residues, where the terminal Cβ points in the opposite direction. We repeated the process of rotating and translating as explained above. Having all 4 β-sheets, we relaxed the backbones using harmonic AtomPair constraints between NH- and CO-groups of adjacent β-strands with the FastRelax mover implemented in RosettaScripts^34^. From these relaxed β-sheets, we then built the remaining ones containing 7 and 6 residues per β-strand by removing 1 or 2 residues per β-strand, respectively.

We combined all pairs of 3-stranded and 4-stranded β-sheets of different or same chiralities restricted to a maximum difference of 1 residue between them – i.e., we made the combination between a 3-stranded β-sheet of 6 residues per strand with the 4-stranded β-sheet of 7 residues but not with that of 8 residues. The combinatorial yielded a total of 28 β-sheet pairs, for which we generated 20,000 open backbones each sampling relative β-sheet-β-sheet twist rotations from -60º to 60º, translations between 10 Å to 12 Å and further transformations along two orthogonal axes from -5 Å to 5 Å. We then carried out loop closure of these structures with the Blueprint Builder^14^ mover in two steps: first designing the three β-hairpins (with canonical AA/BG/EA/GG or BAAGB ABEGO conformations for hairpins with L and R chiralities^15^, respectively), and secondly the three β-arches (with loops ranging between 3 and 6 residues in length, and conformationally unrestricted). For each closed scaffold, we designed 5 different sequences with the FastDesign^35^ mover implemented in RosettaScripts. With this method, we obtained a total of 2,451,516 *de novo* Ig domains; 1,106,651 with positive rotations and 1,344,865 with negative twists. The pool of closed designs represents 88% of all sampled backbones.

We probed the capability of the designed sequences to recapitulate the structures with AlphaFold2. For this, we used the local installation (LocalColabFold: github.com/YoshitakaMo/localcolabfold) of the optimized AlphaFold2 software version (ColabFold) for protein structure predictions^36^. For all protein structure predictions with LocalColabFold, the -use_turbo option was enabled for optimal speed, the number of recycles was kept at 3 (default), the predictions were made in the absence of multiple sequence alignment (--msa_method single_sequence) and relaxed using the Amber force field. We calculated structural similarity metrics (TM-score and RMSDs) for all predicted models against their reference and those with averaged composite scores (TM-score * pTM * pLDDT) >= 0.5 and averaged RMSDs <= 1.25 Å across top 3 AF2 predictions with RMSD standard deviations <= 0.25 Å across all 5 predictions were classified as highly confident and accurate *de novo* immunoglobulins and selected for further characterization.

### Analysis of β-sheet-β-sheet twist rotations

We recomputed twist rotations for all closed β-sandwiches including original designs and redesigns (sequence and structure). First, we defined a vector connecting the center of masses of Cα atoms between the termini residues of β-strands E1, E2 and E5 (for the 3-stranded β-sheet) and β-strands E3, E4, E6 and E7 for the 4-stranded β-sheet. Then, we projected these two vectors onto a perpendicular plane to the connecting vector of both β-sheets (from their center of masses). Finally, we calculated the angle between the projected vectors and used it as an estimation of the twist rotation.

### Sequence and structure redesign using deep learning

For sequence redesign, we used ProteinMPNN and generated 50 new sequences for each scaffold with default parameters (--sampling_temp 0.1, –seed 37, –batch_size 1). For structural validations, each of the 50 sequences were threaded onto their original scaffold using the SimpleThreading mover and relaxed with the FastRelax mover, both implemented in RosettaScripts^34^.

For structure redesign, we first aligned each of the selected scaffolds to their corresponding first AlphaFold2 prediction. Then, for each of the six different loops (3 β-hairpins and 3 β-arches), we computed RMSDs and those with RMSD values > 1.25 Å were selected for inpainting (using RFDesign) new loops with the same length or adding/removing 1 residue. Finally, the generated scaffolds were sorted according to their *inpaint_lddt* and a maximum of 5 models were further relaxed.

For both methods, we performed protein structure predictions with AlphaFold2 in the absence of multiple sequence alignment and using default parameters.

### Design of immunoglobulins with positive rotations using RFDiffusion

We randomly selected 100 highly accurate confident and convergent *de novo* immunoglobulin scaffolds with positive rotations with twists ranging from +11º to +37º, designed by our parametric approach, and used it as a template set. For the *fold conditioning* protocol, we first used the helper script “make_secstruc_adj.py” provided by RFdiffusion to obtain secondary structure and block adjacency information from the template set. We then generated 100,000 backbones by conditioning the monomer fold, with default noise parameters and allowing an insertion of 0 to 3 residues in each loop of the scaffold. For the *motif scaffolding* protocol, we generated 100,000 backbones with positive twist rotations by fixing all residues of each β-strand, except for the first and last one. In this way, we used the core of the Ig framework as motif, allowing the design of all ββ loops with lengths ranging from 3 to 10 residues, which also accounts for β-strand extensions if needed by diffusion.

For both protocols, 5 sequences per backbone were generated with ProteinMPNN using default parameters, and were subsequently threaded onto their original scaffold using the SimpleThreading and FastRelax movers both implemented in RosettaScripts. To verify that these designs had the same 7-stranded Ig topology as the parametric designs, we computed inter-strand distances between E1, E2 and E5 (for the 3-stranded β-sheet) and strands E3, E4, E6 and E7 (for the 4-stranded β-sheet),. We then calculated their twist rotation values as explained above.

For the *in silico* validations, we first performed protein structure predictions with ESMFold and default parameters (model = esm.pretrained.esmfold_v1()) and designs with pLDDT scores > 90 underwent protein structure predictions with AlphaFold2 (using default parameters and without multiple sequence alignment or template information). Finally, we calculated structural similarity metrics (TM-score and RMSDs) for all predicted models against their reference (originally generated scaffold from RFDiffusion) and those with averaged composite scores (TM-score * pTM * pLDDT) >= 0.5 and averaged RMSDs <= 1.25 Å across the top 3 AF2 predictions with RMSD standard deviations <= 0.25 Å across all 5 predictions were classified as highly confident *de novo* immunoglobulins.

## ACKNOWLEDGMENTS

We acknowledge computing resources provided by the Galicia Supercomputing Center (CESGA), and the Red Española de Supercomputación (grants BCV-2021-1-0014 and BCV-2021-3-0010). This research was supported by grants from the Spanish Ministry of Science and Innovation (RYC2018-025295-I, EUR2020-112164 and PID2020-120098GA-I00). J.R.T. was supported by an EMBO postdoctoral fellowship (under grant agreement ALTF 145-2021). L.C. was supported by a predoctoral fellowship from the Spanish Ministry of Science and Innovation (PRE2021-098555).

## DATA AVAILABILITY

Design models of the high-quality immunoglobulin domains are available online (https://zenodo.org/record/8380285). Other data are available from the corresponding authors upon request.

## CODE AVAILABILITY

The Rosetta macromolecular modelling suite (http://www.rosettacommons.org) and the PyRosetta software (https://www.pyrosetta.org) are freely available to academic and non-commercial users. Computational protocols for designing protein structures are available at https://github.com/JorgeRoel/betasandwich.

