## supplementary for "The structural landscape of the immunoglobulin fold by large-scale *de novo* design"

***for***

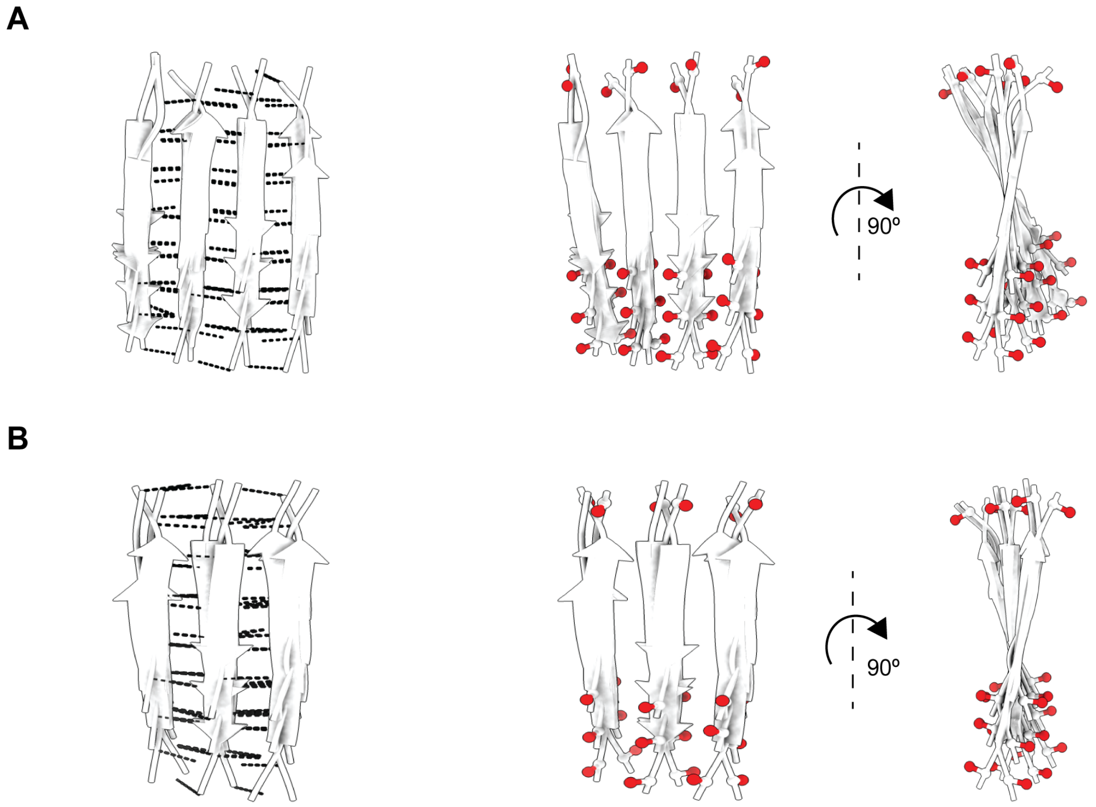


**Figure S1.** Precomputed library of ideal β-sheets formed by 4 (A) or 3 (B) antiparallel β-strands with optimal backbone hydrogen bond pairing, no register shift and sampling the two possible sidechain directionality patterns per β-sheet. The β-sheets sample residue lengths from 6 to 8 amino acids per β-strand. Hydrogen bonds are shown as black dashed lines and Cβ of termini residues as red balls.

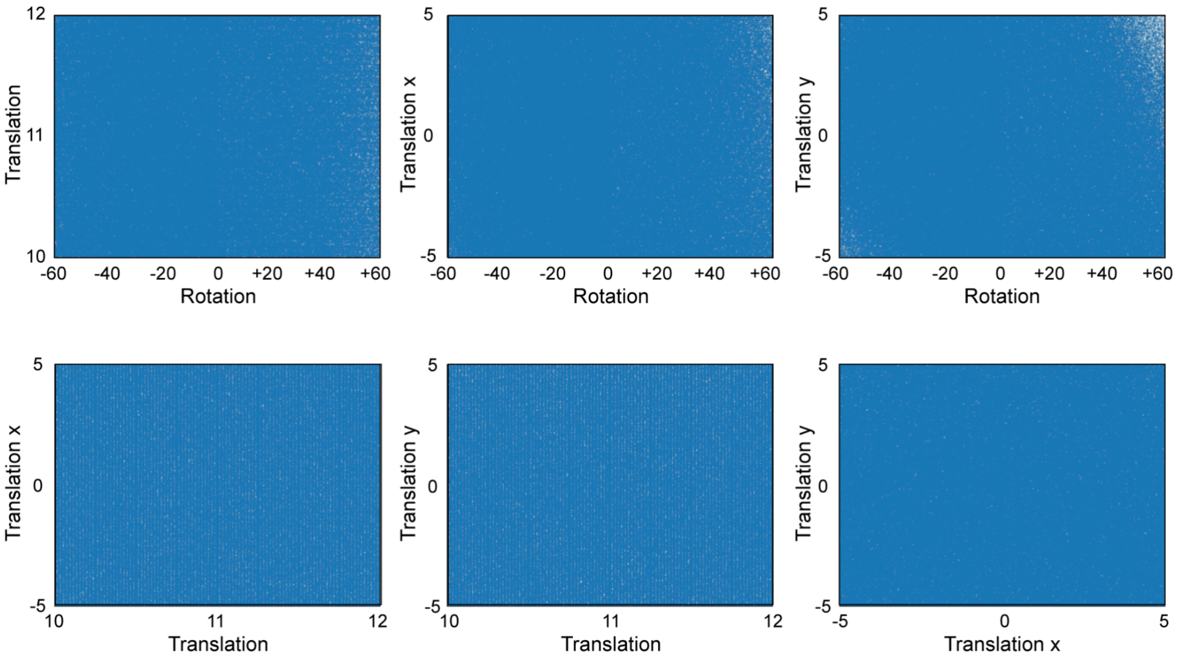


**Figure S2.** Distribution of combinations for the geometrical parameters sampled by our parametric approach on the ~2.5 million *de novo* immunoglobulins. Rotations are expressed in º, while translations in Å.

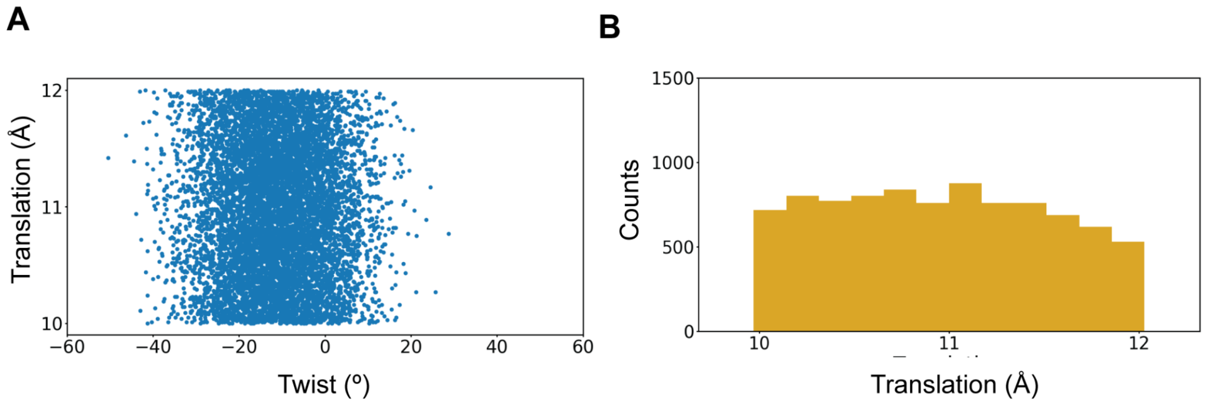


**Figure S3.** Geometrical parameters of the 9,000 high-quality Ig domains designed with the parametric approach. **A**, β-sheet-β-sheet translations and twist rotations were uncorrelated. **B**, Distribution of β-sheet-β-sheet translations.

A

B

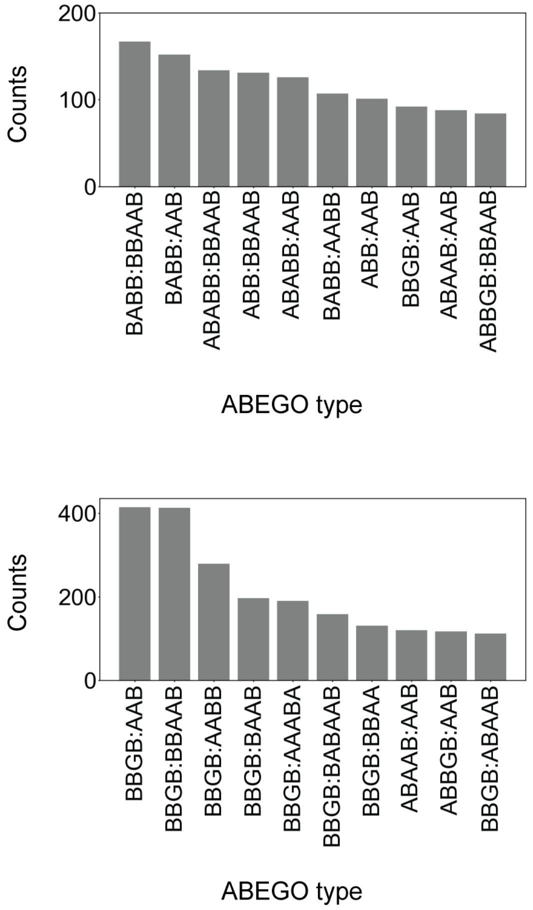


**Figure S4.** Predominant β-arch configurations found in the pool of 45,769 high-quality *de novo* designed immunoglobulins for L2:L5 **(A)** and L4:L5 **(B)**.

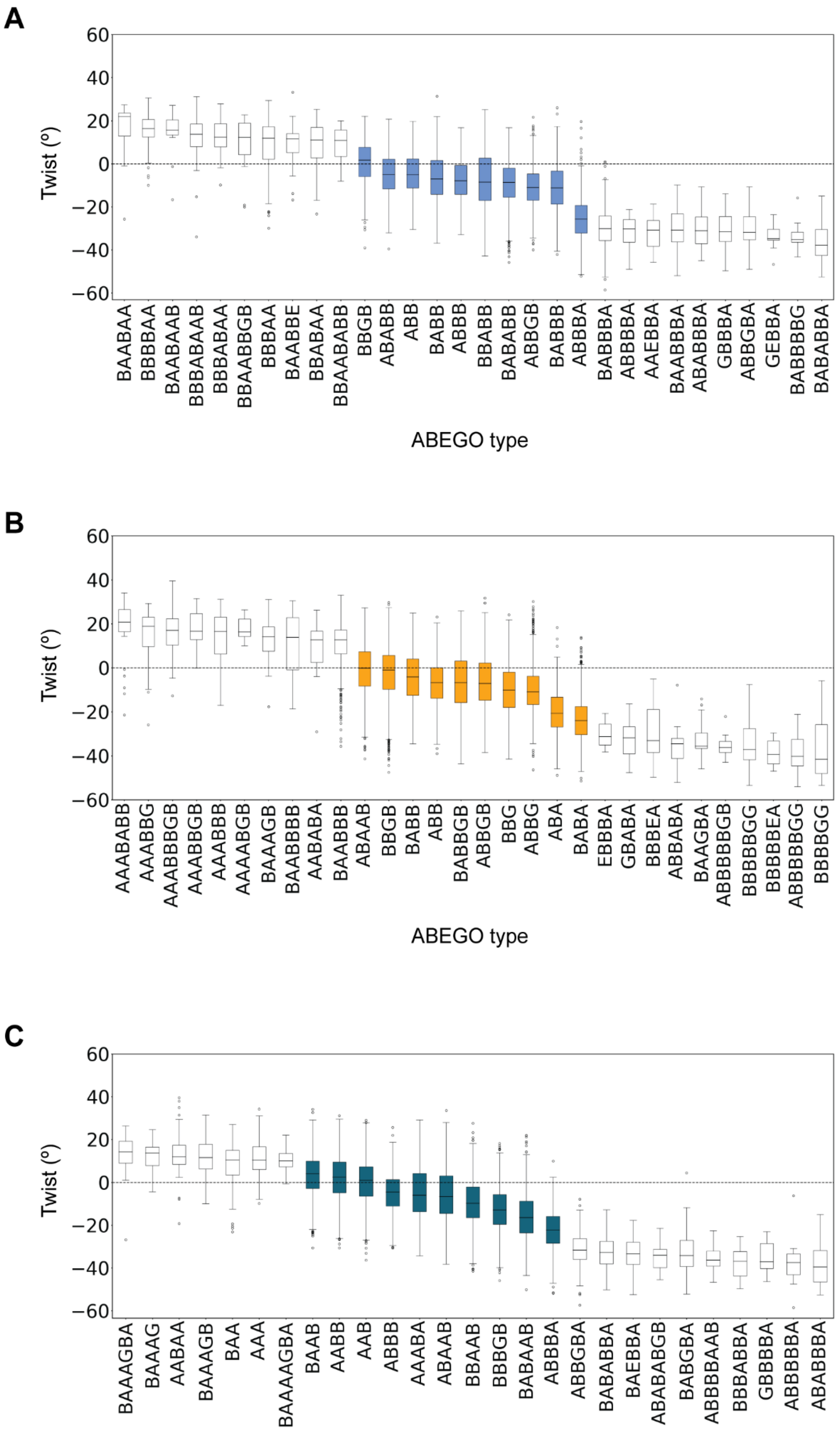


**Figure S5.** Distribution of L2 (**A**), L4 (**B**) and L5 (**C**) β-arches as a function of the immunoglobulin twist. Colored boxes represent the most frequently found ABEGO types.

**A**

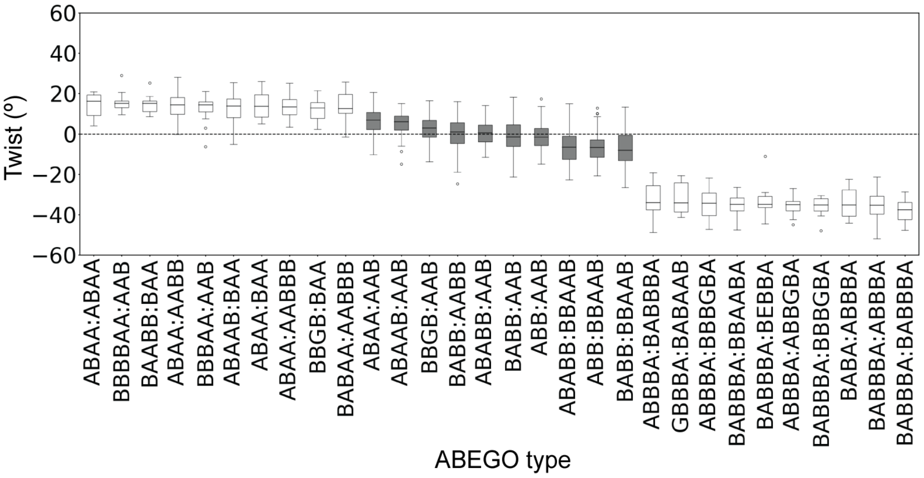


**B**

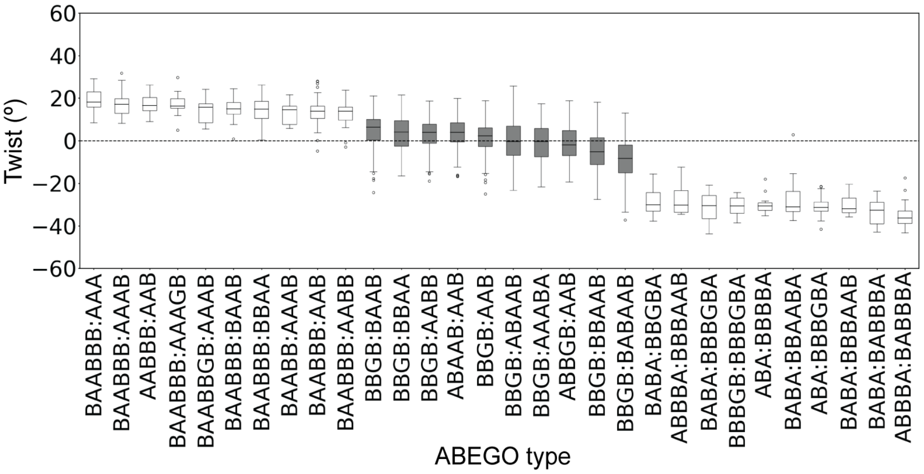


**Figure S6.** Distribution of L2:L5 (**A**) and L4:L5 (**B**) β-arch pairs as a function of the immunoglobulin twist. Gray boxes indicate the most frequently found pairs.

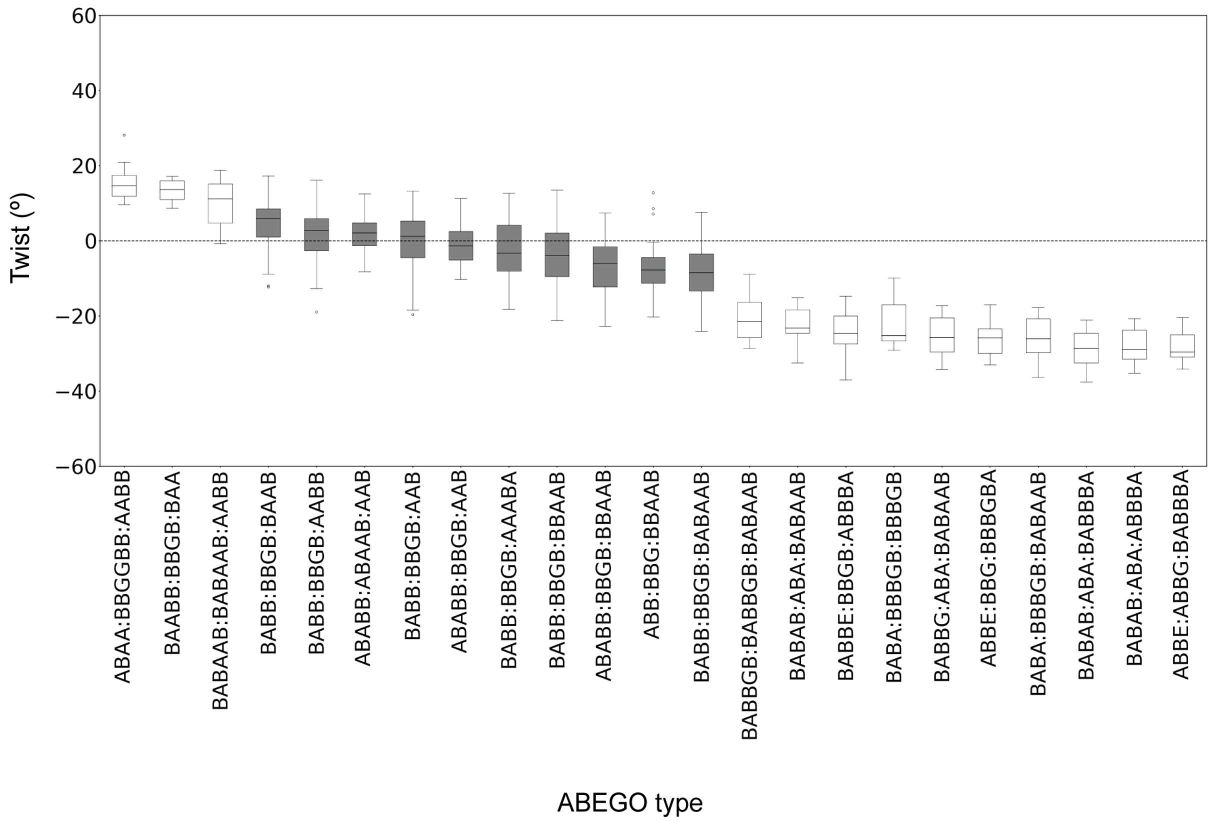


**Figure S7.** Distribution of L2:L4:L5 β-arch trios as a function of the immunoglobulin twist. Gray boxes indicate the most frequently found trios.

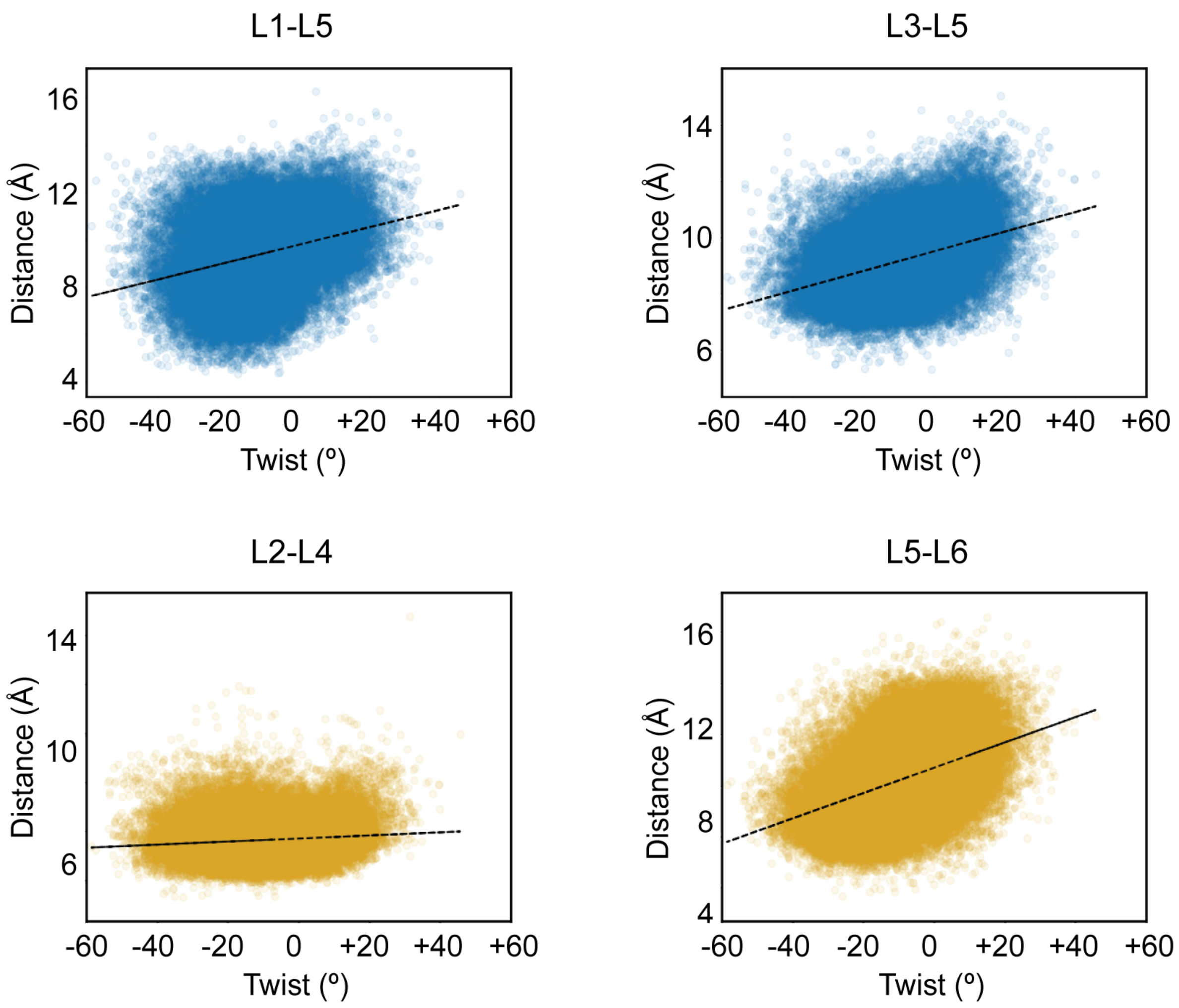


**Figure S8.** Distribution of distances between the center-of-masses of L1:L5, L3:L5, L2:L4 and L5:L6 (y axis) as a function of the β-sheet-β-sheet twist (x axis).

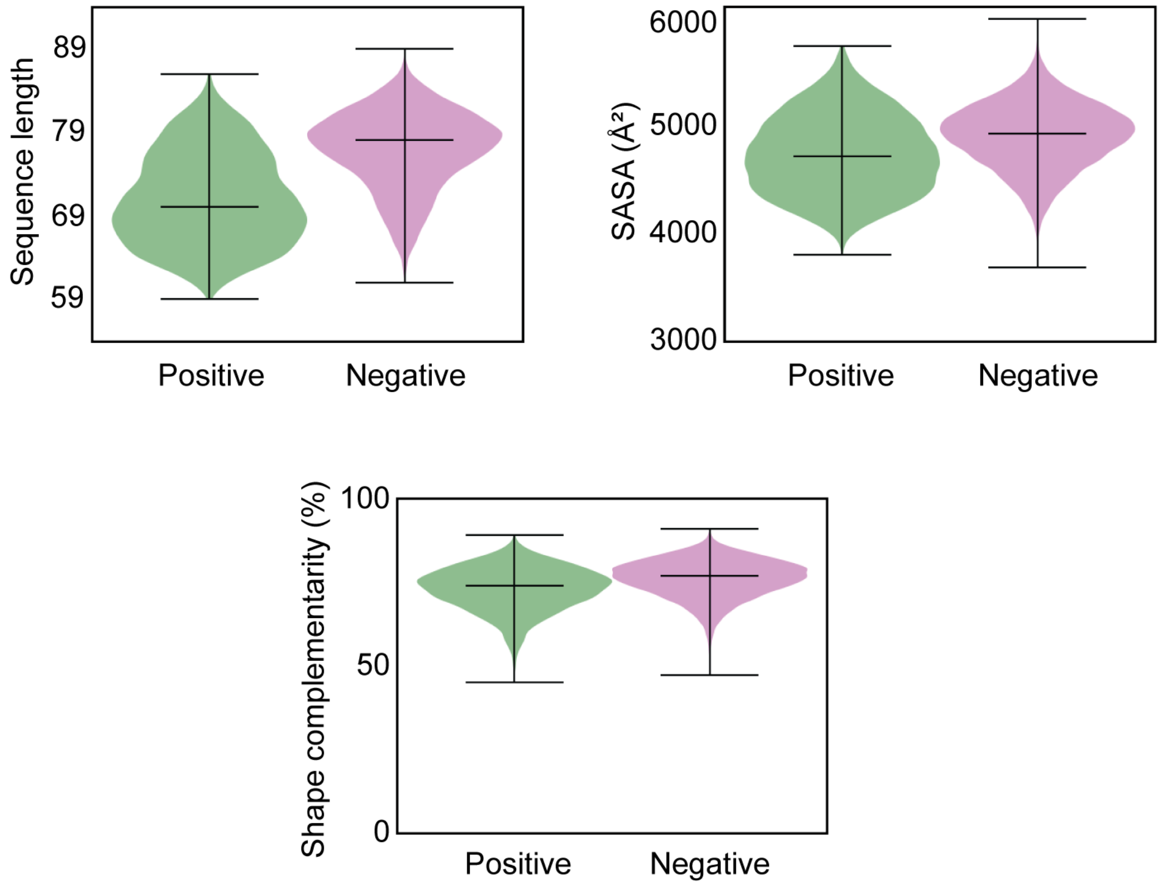


**Figure S9.** Distributions of protein size, solvent accessible surface area (SASA) and the β-sheet-β-sheet shape complementarity for high-quality immunoglobulin domains with positive or negative twist rotations.

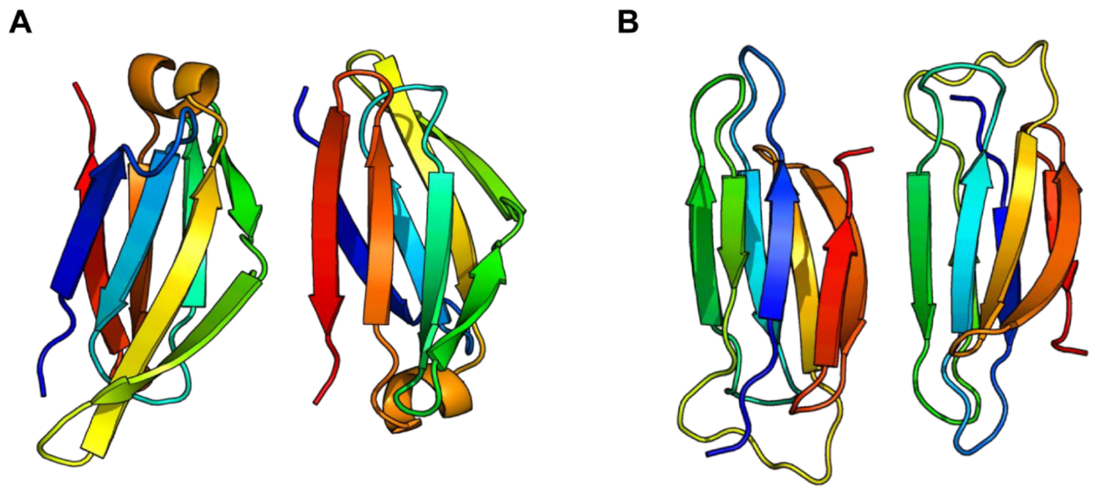


**Figure S10.** Design examples generated with RFDiffusion. **(A)** less regular immunoglobulin domain generated by *fold conditioning* and **(B)** misfolded domains with wrong β-strand pairing and ββ connections. Proteins are colored from N-terminal (blue) to C-terminal (red).

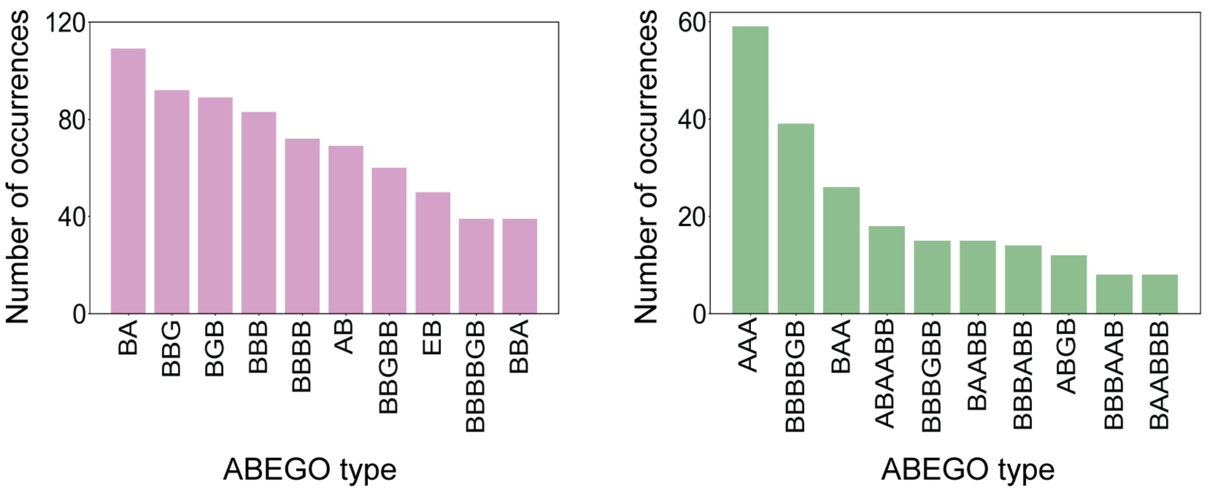


**Figure S11.** Number of occurrences of most frequently found β-arch conformations in *de novo* designed immunoglobulins with negative (left) or positive (right) β-sheet-β-sheet twist rotations as compared to naturally occurring protein structures.
